# Targeted Intestinal Cooling Offers Superior Brain Protection in the Mouse Stroke Model

**DOI:** 10.1101/2025.10.10.681764

**Authors:** Chunli Liu, Yujung Park, Kurt Hu, Yamileck Olivas, Bingren Hu

**Affiliations:** Departments of Emergency Medicine and Neurosciences, University of California San Diego, La Jolla, CA; Department of Medicine, Division of Pulmonary, Critical Care, and Sleep, Medical College of Wisconsin, Milwaukee, WI; Veterans Affairs San Diego Healthcare System, 3350 La Jolla Village Dr, San Diego, CA

**Keywords:** Mice, temperature management, middle cerebral artery (MCA) occlusion (MCAO), hypothermia, intestinal cooling, body surface cooling, histopathology, body weight, nest building activity, pole test, survival rate and mortality

## Abstract

**Background:** Intestinal immune and inflammatory response plays a detrimental role following a stroke. This study aims to evaluate the brain protective efficacies of a novel intestinal cooling (CC) technique relative to the body surface cooling (SC) and the normothermic (NT) condition in a mouse stroke model.

**Methods:** Mice were randomly assigned to CC (n=13), SC (n=15), or NT (n=11) groups. They underwent 60 min of middle cerebral artery occlusion (MCAO) followed by 7-day reperfusion. Both head and intra-colon temperatures were maintained at 37°C for 30 min before, during, and 30 min after MCAO. At 30 min reperfusion, a cooling catheter was placed to maintain intra-colon at 37°C in NT or 12°C in CC. The head temperature was maintained at 37°C in NT and 30°C in CC. In SC, both intra-colon and head temperatures were maintained at 30°C. Cooling lasted 3 hours. Bodyweight, behavioral deficits (nesting and pole test), and survival rate were assessed post-MCAO. At day 7 post-MCAO, mice were perfusion-fixed for histopathological analysis.

**Results:** Post-stroke histopathological brain injury areas and volume were significantly reduced in CC, and appeared reduced though not statistically significant in SC, relative to NT. Compared with NT, body weight, nest building activity, and pole test were all significantly recovered in CC post-MCAO. In SC, only nest building improved significantly, while body weight and pole test showed marginal, nonsignificant trends. Consistent with functional recovery, survival was significantly improved in CC but not in SC, compared with NT.

**Conclusion:** In a murine model, our novel CC technique successfully achieved targeted intestinal cooling while preserving safe upper-body temperatures necessary for normal cardiopulmonary function. Targeted intestinal cooling provides significant benefits superior to SC and NT, including smaller stroke volume, fewer functional deficits, and lower mortality rates, thus supporting the novel concept that the intestines are potential therapeutic targets for stroke management.

**Graphical Abstracts:** 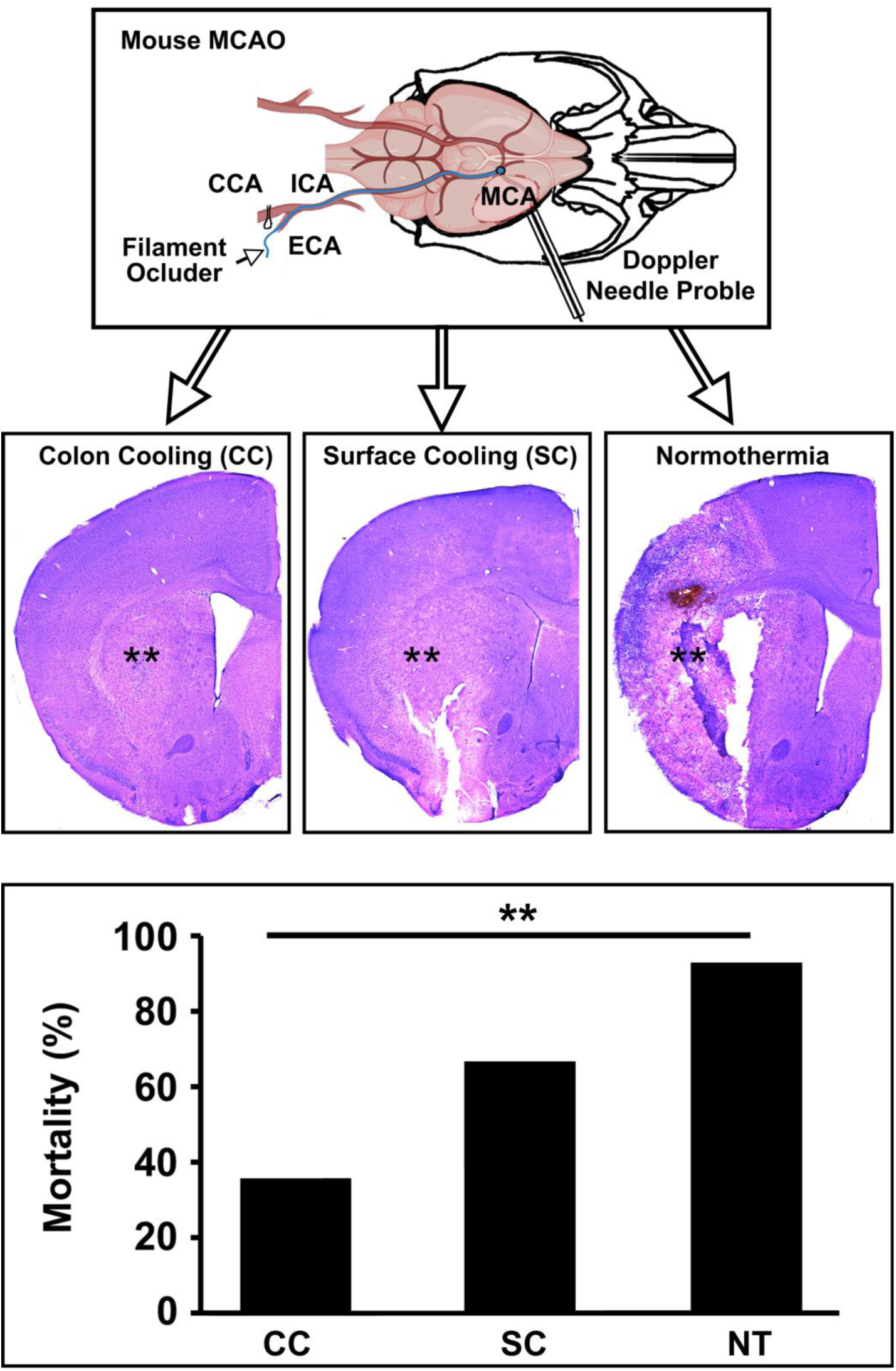

**Highlights:**

- Post-stroke body surface cooling-induced hypothermia demonstrated marginally neuroprotective effects compared to normothermic conditions in the mouse model.
- Targeted deep cooling of the intestine after a stroke resulted in a significantly greater reduction in stroke injury areas and volume, as well as lower mortality rates and fewer functional deficits, compared to body surface cooling-induced hypothermia and normothermic conditions.

## INTRODUCTION

Under normothermic conditions, brain tissue tolerance to ischemic stroke is extremely limited. Brain injury can occur in as little as 5-10 minutes.^1–4^ Currently, hypothermia is the most effective protective treatment for ischemic brain injury.^3,4^ Previous studies have demonstrated that lowering core body temperature to around 10°C offers optimal survival rate and protection in animal hemorrhagic cardiac arrest models.^2,5,6^ However, two major hurdles prevent the use of “deep” (∼18°C) or “profound” (∼10°C) hypothermia in clinical practice. First, rapid cooling of an adult human body is not feasible without extraordinary measures.^3,4^ In addition, core body temperature below 28°C is often associated with sudden cardiopulmonary failure.^7,8^

To investigate solutions to these challenges, the mouse middle cerebral artery occlusion (MCAO) model is often used. These models consistently demonstrate a time-dependent progression of histopathological damage, physical (i.e., body weight) and functional deficits, as well as local and systemic inflammatory responses. The progression includes early infarction in the dense ischemic core (1-12 hours), followed by delayed neuronal death in the penumbra (12-72 hours).^9,10^ After 24 hours, microglia activation, infiltration of various inflammatory cells, and astrogliosis become more apparent in damaged brain areas.^9,10^

The objective of the present study was to employ a mouse stroke model to investigate several key aspects of targeted intestinal deep cooling treatment. We aimed to determine whether a targeted deep colon cooling (CC) technique could provide faster cooling of the intestines and surrounding organs compared to the currently available surface cooling (SC) technique. Additionally, we evaluated the efficacy of the CC, relative to the SC technique, for the management of ischemic stroke. Finally, we assessed the impact of post-MCAO CC and SC on post-stroke brain injury volume, survival rate, as well as physical and functional recovery, compared to the normothermic (NT) condition.

## MATERIALS AND METHODS

A detailed descriptions of Materials and Methods were included in the supplementary materials.

### Ethics Statement

The animal protocols were approved by the Institutional Animal Care and Use Committees (IACUCs) at the University of Maryland, Baltimore, and the University of California, San Diego.

### Animals

Male C57/B6 mice at about 3 months of age were given food and water ad libitum and kept on a 12 h light/dark cycle in climate-controlled housing. They were bred, housed, and cared for following institutional guidelines.

### Mouse MCAO Model

On the day of surgery, the mouse’s body weight, rectal temperature, surgical procedures, and anesthesia status were documented. Aseptic surgical procedures were performed throughout the experiment. Mice were subjected to 60 minutes of MCAO followed by 7 days of reperfusion, as described in our previous study.^11^ Briefly, the middle cerebral artery (MCA) blood flow was monitored through a small incision to attach a needle laser Doppler flowmetry (LDF) probe (MNP100XP, AD Instruments, CO, USA) to the surface of the temporal bone (Supplement Fig. 2A).^12^ A midline incision was made in the neck area, and the right common carotid artery (CCA), external carotid artery (ECA), and internal carotid artery (ICA) were separated. The CCA and ICA were briefly clamped (Supplement Fig. 2A).^12^ A 12 mm sterilized monofilament (6-0, Nylon suture, United States Surgical, Norwalk, CT, USA) with a silicone-coated tip measuring 0.22-0.23 mm in diameter and 2.0 ± 0.2 mm in length was carefully introduced via the ECA into the ICA (Supplement Fig. 2A), but it was not advanced further into the base of the MCA. A temperature probe was inserted into the descending colon to measure the “intra-colon temperature.” Another probe was inserted into the soft palate area of the mouth to measure the head temperature. From this point onward, the LDF probe was attached directly to the surface of the temporal bone without a holder, and MCA blood flow was continuously monitored throughout the entire peri-MCAO or sham surgery periods (Supplement Fig. 2A).^12^ After securing the LDF probe recording, the already inserted monofilament occluder was gently and gradually advanced to the base of the MCA without disturbing the LDF probe recording. The monofilament occluder was confirmed to have reached the base of the MCA when blood flow in the MCA dropped sharply to below 10% of the pre-ischemic level (Supplement Fig. 2B).^12^ After 60 minutes of MCAO, the monofilament occluder was removed to start reperfusion (Supplement Fig. 2B).^12^ After the post-stroke temperature management period of 3-4 hours, isoflurane was discontinued, and all incisions were sutured. The mouse was returned to its home cage for post-stroke care.

### Flowchart and Experimental Group

Based on data from our recent studies using the same mouse MCAO model^11,12^, a power analysis (power 0.80, alpha 0.05, and beta 0.2) showed that more than 10 mice per group were required for the behavioral and histopathological analysis. Flowcharts and timelines are shown in Supplement 2. A total of 39 mice were randomly assigned to three experimental groups: (i) normothermic group (NT) (n=11); (ii) colon cooling (CC) group (n=13); and (iii) surface cooling (SC) group (n=15). The day before surgery to produce the MCAO model, mice underwent physical (body weight and rectal temperature) and behavioral (nesting building activity and pole test) exams. They showed normal physical conditions and behavioral activities. On the day of surgery, mice were anesthetized and subjected to a surgical procedure that induced 60 minutes of MCAO. The CC, SC, and NT temperature managements were performed during 0.5 to 3.5 hours of the reperfusion period. During the post-MCAO period, mice were examined daily for body weight and nest-building activity, and at 1, 3, and 7 days of reperfusion, they underwent the pole test. At 7 days of reperfusion, mice were perfusion-fixed, and brains were sectioned for histopathological analysis (Supplement Fig. 2).

### Temperature Management

A TransRectal Intra-Colon (TRIC) temperature management device consists of an inflow inner polyethylene terephthalate (PET) tube, a PET outer tube, and a series of fittings at the proximal end of the device, which was described in detail in our previous studies.^13–16^ The TRIC device was inserted via the rectum into the descending colon until a resistance was felt at about 3-5 cm of length from the anus according to the technique described in our previous studies (Liu et al., 2021; Arya et al., 2021 and 2023).^13,15,16^ TRIC temperature management began 30 minutes after reperfusion following 60 minutes of MCAO to cool the colon’s inner wall to 13-15°C (CC group) or maintaining it at 37°C (NT group) for 3 hours. During this period, head temperature was maintained as close to 37°C as possible in both groups using a warming blanket and an overhead lamp. Due to deep colon cooling, the head temperature in the CC group could only be maintained at about 30°C during the 3-hour cooling period. After 3 hours of post-MCAO temperature management, rewarming started at a rate of 0.15°C/min in the CC group until the head temperature reached about 30°C, which took roughly an additional 90 minutes. Therefore, the total TRIC temperature management in the CC group lasted approximately 270 minutes post-MCAO. In the SC group, two plastic bags filled with a very thin layer of ice covered the entire animal’s body starting at 30 minutes after reperfusion following 60 minutes of MCAO to keep both the colon’s inner wall and head temperatures around 30°C for the same 270-minute period after MCAO as the CC group. In the NT group, both the intra-colon and head temperatures were maintained at 36-37°C during the same 270-minute post-MCAO period. At the end of temperature management, all catheters were removed, incisions closed, and isoflurane discontinued. Once the mice were stable and breathing spontaneously after stopping anesthesia, they were extubated and returned to their home cages.

### Post-temperature management care

After temperature management, the mice were returned to the home cage. During the first five consecutive days after MCAO, all post-stroke mice received thermal support by placing a water-circulating heating pad (Cat# 50303, IL, USA) set to circulate water at 35-36°C under one-third of the home cage bottom, which maintained the intra-cage temperature at about 25-26°C. Additionally, each mouse received a daily subcutaneous injection of sterile Ringer’s solution, warmed to about 36°C, at a dose of 0.25 ml per 25 grams of body weight for five consecutive days.

### Behavioral outcomes

All behavioral outcome measurements were conducted in a double-blind manner. The following behavioral outcomes were monitored and documented during the post-surgery periods: (i) daily body weight; (ii) daily nest-building activity; (iii) pole test at 1, 3, and 7 days post-MCAO; (iv) daily motor and sensory neurological functional scores; and (v) time of animal death.^11,12^

#### Body weight

Animal body weight was documented on the day before MCAO and daily at about 10 am after MCAO until the endpoint.^11,12^ Animal body weight indicates the animal’s physical recovery.

#### Nest-building activity

Mice were housed individually with standard corn cob-chip bedding but without environmental enrichment items and had unlimited access to food and water throughout the experiments. The nest-building activity was naturally performed in the home cage, as described in our previous study with some modifications.^11^ The nestlet used in this study is a 5 cm square of pressed cotton (Ancare, Bellmore, NY). Only nestlets equal to 2.5 g were used. If a nestlet was heavier than 2.5 g, it was trimmed to that weight. Our pilot tests showed that mice could tear more than 2.5 pieces of 2.5 g pressed cotton nestlets per day. Therefore, each cage was provided with three nestlet pieces at 10 AM. The following day at 10 AM, the amount of torn and untorn nestlet material in each cage was weighed. Untorn pieces were defined as those weighing more than 0.05 g. The percentage of torn versus untorn nestlet material was then calculated. Mice were tested the day before surgery and daily afterward.

#### Pole test

The mouse forelimb strength and ability to grasp and balance on a vertical pole were evaluated. A 90 cm vertical steel pole with a 10 mm diameter was wrapped with tape and placed in a mouse cage for testing. To cushion the floor, two layers of bubble wrap, covered with a blue pad, were used. The mouse was placed in the testing box for 2 minutes to adapt to the new environment. Thereafter, the mouse was positioned head downward on the pole with its forelimbs placed at the 60 cm mark above the ground. The total time for the mouse to walk to the ground floor (defined as all four limbs leaving the pole) was recorded using a stopwatch. Mice were pre-trained three times in the same way before the main test, with 30 seconds rest between trials. Each mouse performed three runs, and the average time to reach the ground was calculated. If the mouse fell immediately from the 60 cm line to the ground, a total time equal to the longest time in the group was given. The mouse had to walk continuously without stopping; if it paused, the test was canceled and repeated, and paused test was not counted. If the mouse slid down from the pole, a time of 25 seconds was recorded. Tests were conducted the day before surgery and on days 1, 3, and 7 after surgery. The apparatus was cleaned with 70% ethanol and clean water after each trial.

#### Animal death time

Subject death time and possible etiology were documented during the day by investigators and during the night (between 1700 and 0930) using the PhenoTyper infrared video device and its software (Noldus, VA, USA). During the night, mice were continuously monitored by video surveillance, and mice showing no breathing or movement for more than 1 hour were considered dead. Death was confirmed the next morning by investigators.

### Quantitative Histopathological Analysis

The size of the brain damage was quantitatively assessed in a double-blind manner using the same method described in our recent publications and detailed in the Supplementary Materials and Methods.^11,12^ Hematoxylin and eosin (H&E) staining was performed on brain sections. Quantitative analysis was performed on every tenth 50 μm-thick H&E-stained section, and seven brain sections between 1.42 mm and −1.69 mm relative to bregma, spaced at regular 0.5 mm intervals, were selected in a consistent manner for the analysis. The stroke damage area was calculated based on the difference between the contralateral normal area and the ipsilateral normal area to account for brain swelling or other post-ischemic changes, using the MBF imaging system and SteroInvestigator software (MBF Biosciences, VA, USA). Stroke damage volume was calculated

### Statistics and analysis

Statistical analysis was performed using KaleidaGraph software (Synergy Software, PA, USA). A two-tailed t-test was used to analyze data from the two experimental groups. One-way ANOVA followed by Tukey’s test was employed for data from more than two groups. The Wilcoxon-Mann-Whitney test (2*1-sided exact) was utilized for analyzing behavioral performance data from two groups. The Kruskal-Wallis test was used to compare behavioral data from three or more groups. The log-rank test was used to analyze the survival rates, while the chi-squared test assessed mortality rates. Data are presented as means ± SEM, and a p-value less than 0.05 was considered statistically significant.

## RESULTS

### Post-MCAO temperature management

Fig. 1 shows the average head and intra-colon temperatures. In the normothermia (NT) group, both head and intra-colon temperatures were maintained within a range of 36-37°C (Fig. 1, A and B, circle). In the surface cooling (SC) group, both head and intra-colon temperatures were maintained at about 30°C (Fig. 1, A and B, square). In the colon cooling (CC) group, the head temperature was kept in the same range as the SC group. In contrast, the intra-colon temperature was maintained as low as possible, i.e., in the 13-20°C range during the 3-hour post-MCAO cooling period (Fig. 1, A and B, triangle). As a result, differences in head temperature were significant only between NT and CC or SC (p<0.01), but not between CC and SC (p>0.05). In comparison, differences in intra-colon temperature were significant not only between NT and CC or SC (p<0.01) but also between CC and SC (p<0.01).

**Figure 1.**
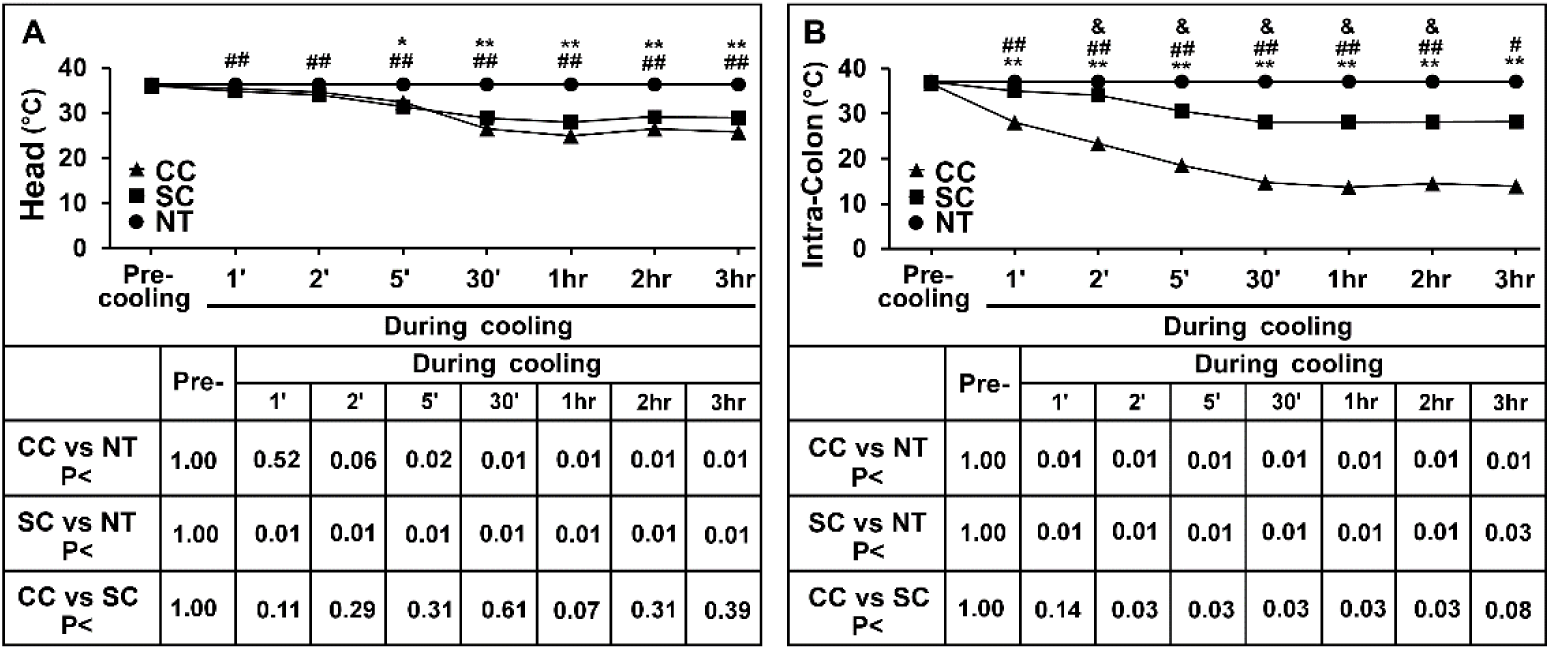
Head (A) and intra-colon (B) temperature during the post-ischemic temperature management period. Colon cooling (CC, n=13), body surface cooling (SC, n=15), and Normothermia (NT, n=11) temperature managements were started at 30 min reperfusion after 60 min MCAO and continued thereafter for 3 hours. Data shown are mean ± SEM, *p<0.05 and **p<0.01 between CC vs. NT; #p<0.05 and ##p<0.01 between SC vs. NT; and &p<0.05 and &&p<0.01 between CC and SC; one-way ANOVA followed by Tukey test.

### Post-MCAO cooling marginally reduces the MCA blood flow

Fig. 2 shows that all mice achieved a relative drop to below 10% of the pre-ischemia baseline in MCA blood flow during the MCAO period, followed by a recovery of MCA blood flow to establish reperfusion. Relative to the peak MCA blood flow at 10-30 minutes of reperfusion, a gradual and significant decrease in MCA blood flow from 10 to 210 minutes of reperfusion was observed in all groups (CC, SC, and NT) (Fig. 2B, p<0.05). The reduction in MCA blood flow was more pronounced in the CC and SC groups compared to the NT group, but the differences did not reach statistical significance (Fig. 2B, p>0.05).

**Figure 2.**
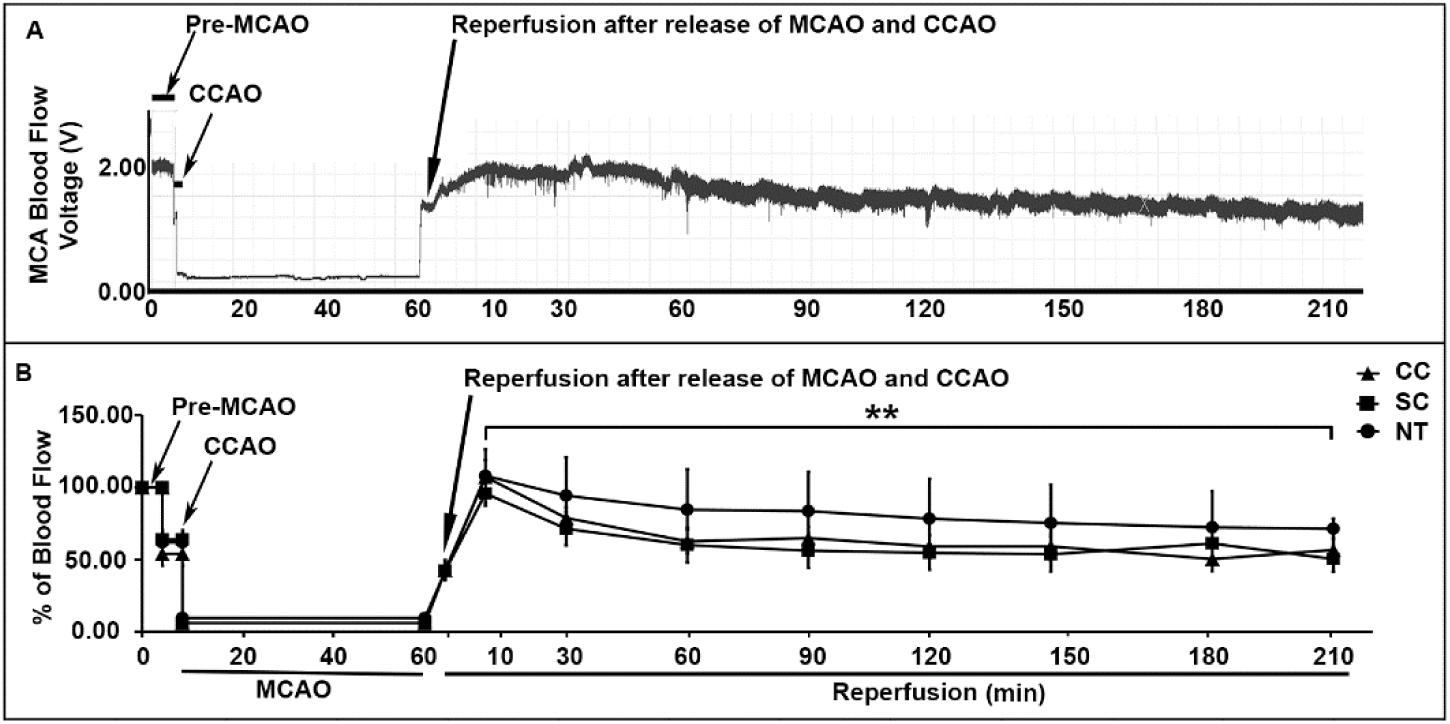
Laser Doppler flowmetry of blood flow in the middle cerebral artery (MCA) territory during peri-MCAO periods. ***A.*** A representative Laser Doppler flowmetry: (i) Pre-MCAO: the MCA blood flow baseline before CCAO and MCAO; (ii) CCA (temporal) occlusion during the MCAO period: MCA blood flow was reduced after common carotid artery occlusion (CCAO); (iii) MCAO period: MCA blood flow was further reduced to the >90% of the pre-MCAO level after CCAO and MCAO with the monofilament; and (iv) Reperfusion: MCA blood flow was resumed close to the pre-ischemia level after release of MCAO and CCAO to start reperfusion. ***B.*** The MCA blood flow in CC (triangle, n=13), SC (square, n=15), and NT (circle, n=11) groups during peri-MCAO periods. The MCA blood flow declined significantly during the period from the 10 min to 210 min reperfusion among CC, SC or NT groups. Data shown are mean ± SEM, **p<0.01 between 10 vs. 210 min reperfusion among all CC, SC or NT groups; no statistically significant differences between CC or SC vs NT groups, one-way ANOVA followed by Tukey test.

### Post-MCAO CC and SC cooling reduces histopathological stroke lesion area and volume

In the CC mice (Fig 3A), brain damage was confined to the dorsal and lateral striatum (−0.15 mm from bregma), while in SC mice (Fig 3B) damage extended to part of the overlying cortical region. In NT mice (Fig 3C), this damage further expanded to nearly the entire dorsal and lateral striatum and cortical region. Seven brain section planes between 1.42 mm and −1.69 mm from bregma, spaced at consistent 0.5 mm intervals, were selected in a consistent manner for quantitative analysis. Fig. 3D shows that stroke areas in five of the seven brain section planes were significantly smaller in CC compared to NT mice (Fig. 3D, **p<0.01; *p<0.05). Stroke volume was also significantly smaller in CC relative to NT brains (Fig. 3E, *p<0.05). Although both stroke areas and volume appeared smaller in CC compared to SC, and in SC compared to NT brains, these differences did not reach statistical significance (Fig. 3E, p>0.05).

**Figure 3.**
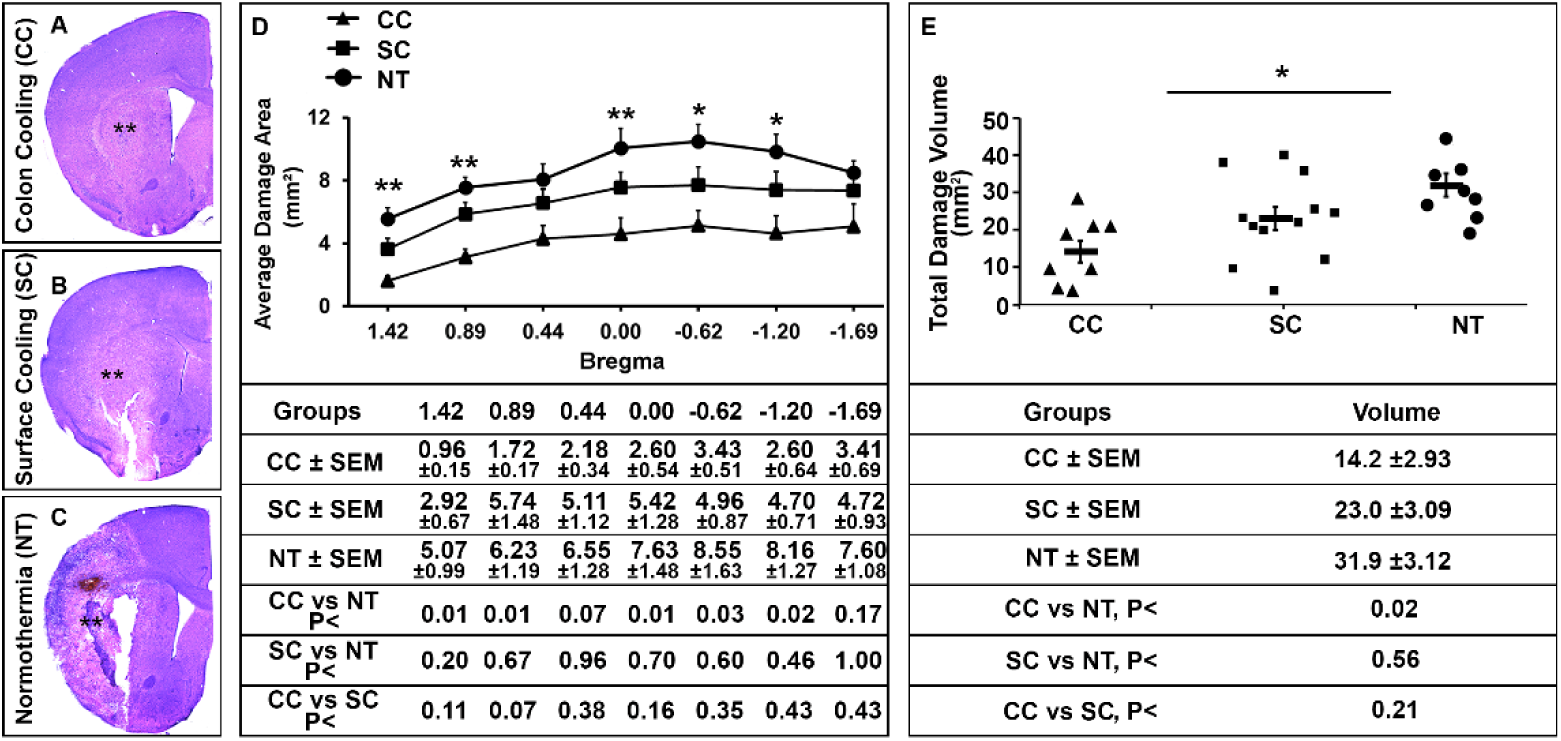
Histopathological brain injury. **A-C**: Representative coronal sections of mouse brains at the striatal and hippocampal levels post-MCAO from CC (A), SC (B) and NT (C) mice. Drawing area indicates stroke damage. **D and E:** Average damage area relative to bregma and stroke volume (E) among CC (triangle), SC (square), and NT (circle) groups. Seven brain sections between 1.42 mm and −1.69 mm to bregma with a regular interval were selected in a consistent manner for the analysis. Ipsilateral damage area was calculated based on a difference between the contralateral and ipsilateral normal areas. The stroke damage volume was calculated by summing the damage areas and section thickness across all sections. Data shown are mean ± SEM. *p<0.05 and **p<0.01, CC vs. NT. Kruskal-Wallis test followed by Bonferroni correction was used for the statistical analysis.

### Post-MCAO CC improves post-MCAO physical and functional recovery

Mouse body weights, used as a sign of physical recovery and measured daily after surgery through day 7 post-MCAO, were consistently lower during the first 2-3 days post-surgery for both post-MCAO and sham-operated mice, with a gradual recovery observed in the following days. The extent of body weight recovery was significantly greater in the CC group compared to the NT group on days 6 and 7 after MCAO (Fig 4A). Although better body weight recovery was also observed in the CC group compared to the SC group, and in the SC group compared to the NT group, these differences did not reach statistical significance (Fig. 3E, p=0.33 and p=0.09, respectively).

**Figure 4.**
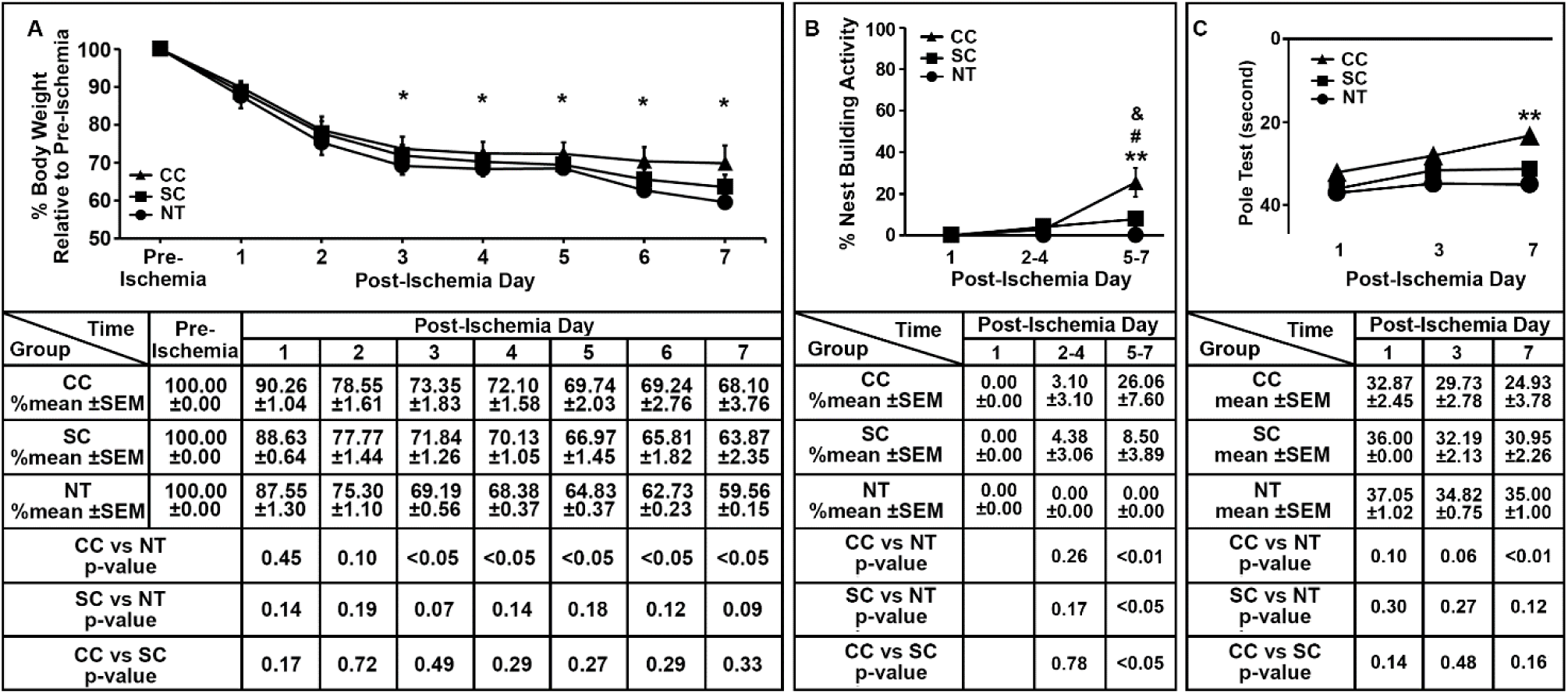
Mouse body weight recovery (**A**) nest-building activity (**B**), and pole test (**C**) after MCAO. Mice were subjected to 40 min of MCAO followed by 7 days of reperfusion, respectively. Body weight, nest-building activity, and pole test were quantified among colin cooling (CC), body surface cooling (SC) and normothermia (NT) groups. Body weight was calculated as percentage of the pre-ischemia body weight. Data shown are mean ± SEM, \**p*<0.05 and **p<0.01, CC vs. NT; #p<0.05, SC vs. NT; &p<0.05 CC vs. SC, Kruskal-Wallis test followed by Bonferroni correction was used for the statistical analysis.

Nest-building activity was nearly abolished in the NT group but showed significant recovery in both the CC (p<0.01) and SC (p<0.05) groups during the 5-7 days after MCAO (Fig. 4B). The extent of recovery was significantly better in the CC group compared to the SC group (Fig. 4B, p<0.05).

The vertical pole was used to assess motor function (Fig. 4C), showing significantly improved performance in the CC group compared to the NT group at 7 days post-MCAO (Fig. 4C, p<0.01), while the improvement in the SC group was not statistically significant (Fig. 4C, p=0.12).

### Survival rate and mortality

At the 7-day post-MCAO endpoint, the CC group showed a significantly higher survival rate compared to the NT group (Fig. 5A, CC vs NT, p<0.05). The SC group did not show a statistically significant increase in survival rate compared to the NT group (Fig. 5A, SC vs NT, p=0.14). Similarly, the mortality rate was significantly lower in the CC group compared to the NT group (Fig. 5B, CC vs NT, p<0.05). The mortality rate in the SC group was between those of the CC and NT groups, but the differences were not statistically significant when compared to either group (Fig. 5B, SC vs NT, p=0.09; CC vs. SC, p=0.28).

**Figure 5.**
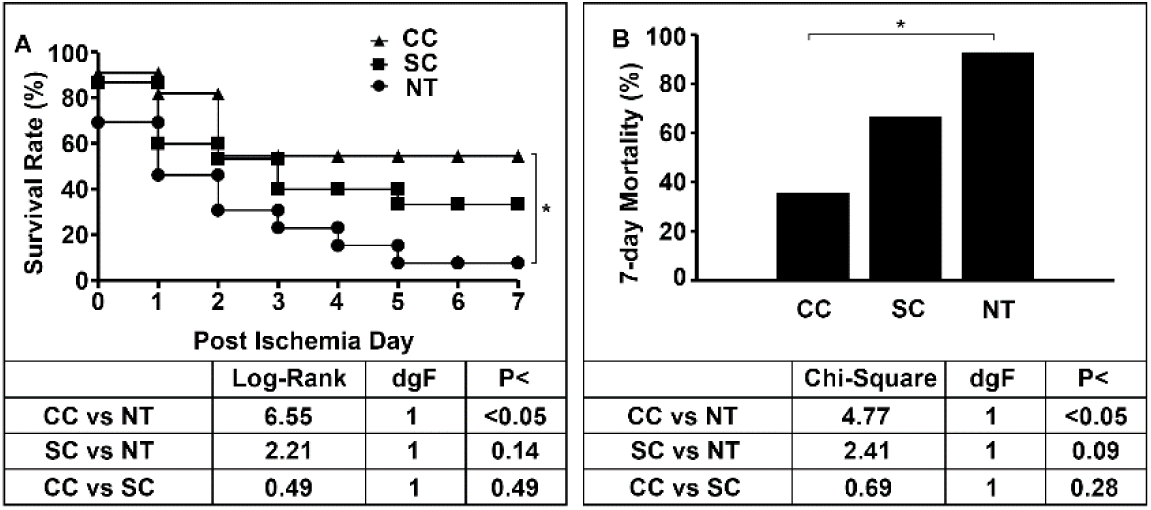
Survival rate (A) and Mortality (B) after 60 min MCAO among colon cooling (CC, triangle, n=13), body surface cooling (SC, square, n=15) and normothermia (NT, circle, n=11) groups. Log Rank test was used for the statistical analysis of Kaplan-Miner survival rate, and Pearson Chi Square analysis was used for the statistical analysis of the mortality, *p<0.05, CC vs. NT.

## DISCUSSION

This study employed a consistent mouse stroke model to explore the impact of intestinal cooling on post-stroke brain injury. Compared to body surface cooling (SC) and normothermic (NT) conditions, post-stroke colon cooling (CC) demonstrated significantly better protection against post-stroke injury. By the 7-day post-MCAO endpoint, the CC group exhibited notable improvements over the NT group in several key areas: reduced stroke volume, faster bodyweight recovery, improved nest-building activity and pole test performance, increased survival rate, and lower mortality. While the SC group showed some marginal improvements in physical, functional, histopathological, and survival outcomes compared to the NT group, these improvements were generally smaller than those seen in the CC group. These findings suggest that targeted intestinal cooling after MCAO provides greater brain protection against stroke injury than systemic body surface cooling-induced hypothermia.

### Rigor and Reproducibility

A significant challenge for using the rodent intraluminal monofilament MCAO model in stroke research comes from a considerable inconsistency in reported infarct volumes up to approximately fivefold.^11,12,17–19^ We attribute these inconsistencies, in part, to inadequate control and confirmation of ischemia and reperfusion. To address these challenges, we continuously monitored the MCA blood flow in every MCAO mouse throughout the peri-MCAO period. Successful occlusion was determined as a sharp drop of the MCA blood flow to less than 10% of the pre-ischemic level (Fig. 2A). At the same time, reperfusion was confirmed by increased in the MCA blood flow when the monofilament was removed (Fig. 2A). This approach strengthens the validity of our finding providing a higher degree of consistency among treatment groups.

### Intestinal cooling may be an improved therapeutic hypothermia modality

The effectiveness of hypothermic protection in ischemic conditions is time dependent, with optimal outcomes achieved when cooling is initiated immediately before or during ischemia.^4,20–23^ While profound hypothermia (down to 10°C) provides maximum protection in lethal hemorrhagic cardiac arrest^5,6,24^, such rapid cooling in humans presents challenges and cardiopulmonary risks.^7,8,25,26^ Furthermore, current targeted temperature management methods like external skin surface cooling and endovascular cooling have limitations in effectiveness and speed due to various reasons.^20,25–27^ Consequently, these methods often fail to achieve targeted temperatures within the critical therapeutic window for conditions such as stroke, neonatal hypoxia-ischemia, post-cardiac arrest, and lethal hemorrhage.^13–16,26^ Lamb et al. developed a method involving wrapping the inferior vena cava and descending aorta with an implanted closed-loop cooling system in the abdominal cavity to induce rapid therapeutic hypothermia in rats.^27^ While this proof-of-concept approach allows faster and more controllable cooling, it is invasive.^27^ A previous study demonstrated that rectal cooling, maintaining tympanic temperature at 33–35°C for 12 hours, significantly reduced hippocampal CA1 neuron death; however, no damage volume or functional outcome assessments were included, as rats were sacrificed at the 24-hour endpoint in that study.^28^

Our study explored the use of non-invasive intra-colonic cooling as an alternative method. Based on previous studies, intra-colon cooling can rapidly cool the abdominal temperature from 37°C to 10°C within 10 minutes while keeping the upper body temperature at around 30°C.^13–16,29^ The effectiveness of this approach likely comes from two key factors: (i) the large volume of circulating blood in the abdominal organs, which are many times larger than that in the skin and subcutaneous fat, and (ii) the use of the abdominal cavity as a natural cooler that efficiently distributes chilled blood to other parts of the body.^13–16,30^ Notably, our previous studies have consistently shown no detectable damage or functional impairment to the intestines or other organs, including the liver, kidney, heart, lung, brain, and spinal cord.^13–16^ Our current study evaluated over a 7-day period, and compared to SC, post-MCAO CC resulted in a greater reduction in stroke volume, improved functional recovery, and a significant decrease in post-MCAO mortality rates in the mouse model.

### Intestinal cooling offers meaningful benefits in the preclinical stroke model

The most appropriate primary outcome measures for animal stroke models are functional outcomes (death or impairment), as recommended by the STAIR panel.^31^ Death and disability rates are always key primary outcome measures in human stroke trials.^31^ However, many of the functional tests currently used in animal stroke studies do not assess animals’ natural behaviors; instead, animals are either “forced” to perform a task (e.g., pole test) during daylight hours, designed by investigators. These tasks are useful for testing functional deficits but may differ from natural benchmarks, such as death and disability.^31^ For that reason, using natural and objective behavioral measures related to death and disability is likely a better analog to human stroke clinical trials.

This study evaluated “natural and objective” recoveries of body weight and nesting building activity as outcome measures among the CC, SC, and NT groups. Both body weight and nest building activity recovered significantly in CC and, to some extent, significantly in SC, compared to the NT group. This study, along with our previous research, has shown that body weight recovery reflects the physical condition of animals post-MCAO and is useful for determining mortality in mouse stroke models.^11^ Consistent with the present findings, previous studies indicate that the degree of body weight loss closely correlates with mortality rates in the MCAO model. Collectively, these studies suggest that body weight recovery is a reliable and straightforward “natural and objective” outcome measure for evaluating brain protective interventions in mouse models.

Both male and female small rodents build nests daily.^32^ Regular nest-building activity requires the coordination of multiple sensorimotor and cognitive functions, indicating mouse activity and well-being.^11^ Nest-building occurs autonomously in the home cage without investigator encouragement.^11^ In this study, nest-building activity was significantly recovered in both the CC and SC groups compared to the NT group, but the recovery was notably better in the CC group than in the SC group. Consistently, previous studies show that the degree of nest-building recovery closely reflects the severity of brain damage.^11^ These studies suggest that nest-building activity may serve as a better “natural and objective” outcome measure than body weight recovery for testing brain protective interventions in mouse models.

Compared to body weight and nest-building activity recovery, “forced” pole test appeared less sensitive as an outcome measure because this test could not consistently show differences over time between the CC and SC groups (see Fig. 4).

### Mechanisms of intestinal cooling for brain protection after MCAO

This study showed that intra-colon cooling after MCAO provides better protection than body surface cooling. While the exact mechanisms behind this improved protection need more investigation, reducing the gastrointestinal (GI) inflammatory response likely plays a key role. Severe stroke patients are prone to develop conventional GI complications and likely GI-mediated immune dysregulation.^33–36^ The GI system contains over 70% of the body’s immune cells and has the largest population of macrophages. Previous research has observed the migration of intestinal macrophages and γδ T-cells to the brain following mouse MCAO.^37–39^ Our earlier study demonstrated that intra-colon cooling prevented intestinal inflammation and helped shift from harmful pro-inflammatory to healing anti-inflammatory responses after fatal hemorrhage in rodents.^15^ These results suggest that cooling the intestine could help reduce the negative effects of the GI-associated immune response after stroke.

### Limitations of the study

Age and sex are important variables in stroke brain injury, but this study only used young healthy male mice and assessed outcomes 7 days after MCAO. Future research should explore how sex and age influence the effects of intra-colon cooling on stroke results. Additionally, the long-term impacts beyond 7 days and the mechanisms behind the increased protection need further investigation. These topics, which are currently being studied in our laboratories, are essential for fully understanding the potential of this technique and for translating it into clinical applications for a wider range of stroke patients.

## Acknowledgments

Kurt Hu, MD, Yujung Park, MD, and Chunli Liu, MD contributed to the study design, animal model, data collection and interpretation, and writing of the manuscript. Yamileck Olivas MD, Chun Mun Loke, BS, Sungeong Lee, BS, and Jian Liang, MD, PhD contributed to the animal model and preparation of graphs. Bingren Hu, MD, PhD, designed the study and wrote the original manuscript draft.

## Sources of Funding

This work was supported in part by National Institutes of Health (NIH) grants: NS097875, NS102815, NS129553, NS130557; and NS134895 to BRH, as well as supported in part by Veteran Affair Merit grant: Award # BX005814 to BRH from the United States (U.S.) Department of Veterans Affairs Biomedical Laboratory Research and Development Service.

## Disclaimer

The contents do not represent the views of the U.S. Department of Veterans Affairs or the United States Government.

## Ethical approval

This article does not contain any studies with human subjects. All the experimental procedures involving the use of animals were approved by the Animal Use and Care Committee at the University of Maryland School of Medicine.

## Declaration of interest

None.

## Declaration of generative AI in scientific writing

None

The authors have stated the sex definition and addressed the sex biovariable as a limitation to the research’s generalizability in Discussion.

## Notes

### Competing Interest Statement

The authors have declared no competing interest.

